# The *Escherichia coli* CpxAR system does not sense surface contact

**DOI:** 10.1101/233999

**Authors:** Tom Eric Pieter Kimkes, Matthias Heinemann

**Author notes:** Correspondence should be addressed to M.H.

## Abstract

For proper biofilm formation, bacteria must have mechanisms in place to sense adhesion to surfaces. In *Escherichia coli*, the CpxAR and RcsCDB systems have been reported to sense surfaces. The CpxAR system is widely considered to be responsible for sensing attachment, to specifically hydrophobic surfaces. Here, using both single-cell and population-level analyses, we confirm RcsCDB activation upon surface contact, but find that the CpxAR system is not activated, in contrast to what had earlier been reported. Thus, the role of CpxAR in surface sensing and initiation of biofilm formation needs to be reconsidered.

**Significance statement:** *E. coli* is capable of forming medically problematic biofilms, which are surface-associated microbial communities, protected by an exopolymeric matrix and with increased antibiotic tolerance. How these bacteria sense physical contact with a surface, which may lead to initiation of the biofilm formation process, is largely elusive. The signal transduction systems CpxAR and RcsCDB have previously been found to activate upon surface contact. Here, we confirm that RcsCDB is a surface sensing system, immediately responding to attachment. In contrast, using two different experimental approaches, we found that the CpxAR system does not perceive surface contact. Thus, contrary to the current view, the CpxAR system does not play a signaling role in the first step of biofilm initiation.

## Introduction

To ensure that the biofilm formation process is initiated only under proper conditions, bacteria need to sense that they are in contact with a surface. Despite the importance of biofilms, both in disease and in technical systems, it turns out that even in the model organism *Escherichia coli* the process of surface sensing is still largely elusive (1–6). There is evidence that in *E. coli* surface contact can be sensed with cell appendages, such as flagella and pili (7–12), but also via the two-component systems RcsCDB, rapidly activated upon contact to hydrophilic surfaces (13), and CpxAR, responding to hydrophobic surfaces (14).

The Cpx system consists of an inner membrane-localized histidine kinase, CpxA, and the response regulator CpxR. Depending on the presence of inducing signals, CpxA can act either as a kinase or as a phosphatase on CpxR (15). While the precise molecular mechanism leading to activation remains to be solved, several inducing cues were found, including extracellular copper (16), osmolarity (17), pH (18, 19), envelope stress (20–23) and, as reported, surface attachment (14). The transcription factor CpxR, in its phosphorylated form, regulates expression of a large number of genes, including a few biofilm-related genes (24).

With regard to induction by surface attachment, expression of CpxR-controlled genes has been reported to increase threefold within an hour after bacteria adhered to hydrophobic glass beads (14). In addition to CpxA and CpxR, also the outer membrane lipoprotein NlpE was suggested to be required for sensing hydrophobic surface contact, and these three proteins were also needed for stable adhesion to hydrophobic surfaces (14). A later study by Shimizu and coworkers (25), using a similar experimental approach, reported surface sensing by CpxAR in a pathogenic *E. coli* strain. Because of the huge biofilm-related problems in both medical and technical areas (26–29), and the currently limited understanding about the initial sensing of surface contact, we aimed at further investigating the CpxAR system with single-cell analyses employing fluorescence microscopy and microfluidics.

Here, while we could confirm that RcsCDB is highly responsive to growth on a surface, we could not confirm the earlier reported response of CpxAR to surface attachment, neither with novel single-cell analyses, nor with the population-level experiments as originally done and reported (14). Our results indicate that RcsCDB, but not CpxAR, is activated upon attachment. Thus, the role of *E. coli’s* Cpx system as a surface sensing system, as widely assumed (2, 4, 5, 30–33), needs to be reconsidered.

## Results

To investigate the single-cell response of *E. coli* to surface contact, we used microfluidics and microscopy. Specifically, bacteria were transferred from an exponential phase culture in M9 glucose medium to the microfluidic device, where they were brought in contact with the surface of the cover glass by placing a polyacrylamide gel pad on top of the cells (Figure 1a). To ensure otherwise constant conditions (apart from the surface contact), the gel pad had been equilibrated with spent medium, which was also constantly perfused over the polyacrylamide pad during the whole experiment.

**Figure 1:**
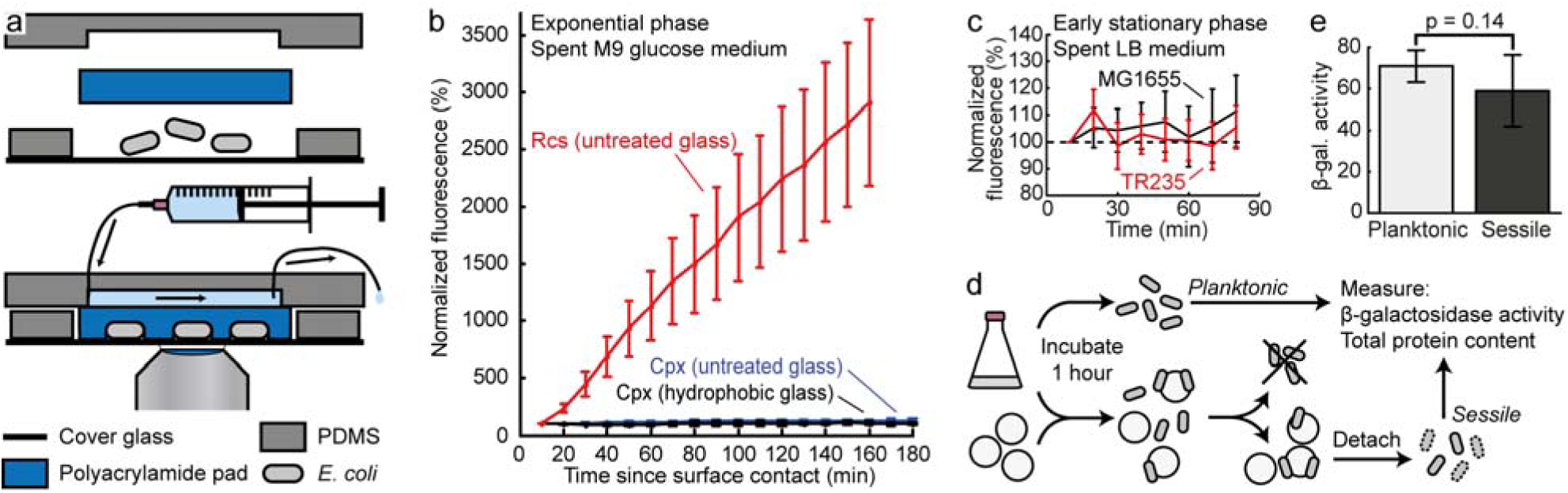
(a) Overview of the microfluidic setup used for the microscopic observation of fluorescence intensity in single surface-attached cells. (b) Comparison of GFP expression from the Rcs-regulated *rcsA* promoter (red; n = 46; 2 independent experiments) and the Cpx-regulated *yebE* promoter (blue; n = 23; 2 independent experiments) on untreated cover glasses, and the *yebE* promoter on hydrophobic cover glasses (black; n = 26; 2 independent experiments) in the microfluidic device with flow of spent M9 glucose medium. The fluorescence intensity of each cell at the first time point is set to 100%. Error bars show 95% confidence intervals. (c) Fluorescence intensity in surface-attached MG1655 + pPyebE-*gfp* (black; n = 60; 3 independent experiments) and TR235 + pPyebE-*gfp* (red; n = 40; 2 independent experiments), grown in LB medium to an OD_600_ of 2 before introduction into the microfluidic system, where there was flow of spent LB medium. The fluorescence intensity of each cell at the first time point is set to 100%. Error bars show 95% confidence intervals. (d) Overview of the population-level assay. *E. coli* TR235 from early stationary phase LB culture were incubated with or without hydrophobic glass beads for 1 h. Unattached cells in the sample with beads were removed and discarded. Attached cells were detached by vortexing in the presence of SDS, which causes the bacteria to lyse. For both the detached sessile cells and the planktonic control, the β-galactosidase activity and total protein content were determined. (e) Comparison of β-galactosidase activity in planktonic and sessile TR235 (MC4100 λRS88[*cpxR*-*lacZ*]). Planktonic: bacteria incubated without beads for 1 h. Sessile: Bacteria that were incubated with hydrophobic beads for 1 h and that had attached. The activity was normalized to total protein content as determined from silver-stained polyacrylamide gels. The values are the mean with 95% confidence intervals (n ≥ 6). The p-value was obtained from a Wilcoxon–Mann–Whitney test.

To confirm that immobilization in the microfluidic setup is perceived as surface contact, *E. coli* carrying a fluorescent transcriptional reporter, controlled by the RcsAB-regulated *rcsA* promoter (13, 34), were transferred to the microfluidic device, and the cells were observed by time-lapse fluorescence microscopy. The cells rapidly became highly fluorescent (Figure 1b). When the *rcsB* gene was deleted, the large increase in fluorescence was no longer observed upon surface contact (Supplementary figure S1a). Thus, the previously reported surface sensing by the Rcs system (13) was also observed in our microfluidic setup, showing that the system can be used to investigate the response of single cells to surface contact.

Towards investigating the surface response of CpxAR, we first tested the functionality of the respective reporter. Specifically, we tested induction of the two-component system by copper, a known activating signal (16). Here, we observed a rapid increase in fluorescence in cells carrying a fluorescent transcriptional reporter under control of the CpxR-regulated *yebE* promoter (16, 24, 35) (Supplementary figure S1b). To exclude a global effect of copper as the cause of the induction, we also tested the reporter for the Rcs system and found that it was not induced (Supplementary figure S1b). Thus, the transcriptional reporter for CpxAR is functional.

To test whether the Cpx system responds to surface contact, similarly as the Rcs system, *E. coli* carrying the reporter plasmid were immobilized in the microfluidic device and followed in time-lapse by fluorescence microscopy. Here, we found that the fluorescence intensity remained unchanged after surface attachment (Figure 1b). As the previous report, where the Cpx system was suggested to respond to surface contact with a threefold induction after one hour (14), had used hydrophobic surfaces, we next performed the same experiment with a cover glass that was rendered hydrophobic. Also here, even though we used the same hydrophobic dimethyldichlorosilane coating as previously (14) used, the fluorescence intensity of attached cells was unaltered (Figure 1b). These results, where we could not find activation of the CpxR system, neither on untreated cover glasses nor on hydrophobic glasses, cast doubts on the earlier reported CpxR surface activation.

As so far, we used glucose minimal medium and exponentially growing cells, we adjusted the growth conditions to mimic those applied by Otto and Silhavy (14): the cells were grown until early stationary phase in LB medium before we introduced them into the microfluidic device. We used a hydrophobic cover glass and the flow of medium over the polyacrylamide pad was spent LB to mimic the conditions in the earlier performed experiments. Also under these conditions, the fluorescence intensity of the bacteria remained unchanged (Figure 1c). Also, experiments with the *E. coli* MC4100 background (TR235, kindly provided by T.J. Silhavy) as earlier used, and transformed with the reporter plasmid, yielded no response (Figure 1c), excluding strain-to-strain differences as the cause. Thus, on the single-cell level we could not find any activation of the Cpx system by surface sensing.

To determine whether the negative results are caused by the different experimental setup, we repeated the original experiments that established CpxAR as a surface sensing system (14), with the same TR235 *E. coli* strain. Specifically, we incubated early stationary phase cells in the presence of hydrophobic glass beads for 1 h, then removed and discarded unattached cells, detached sessile bacteria by vortexing in buffer containing SDS, permeabilized them with SDS and chloroform, and carried out β-galactosidase assays, for which bacteria incubated without beads were used as the planktonic control (Figure 1d). Instead of normalization to optical density of the bacterial sample (as in the original publication), we normalized the β-galactosidase activity to total protein content, as determined from a silver-stained polyacrylamide gel. We altered the procedure, because we found that the removal of attached bacteria from the hydrophobic glass beads causes cell lysis if carried out as described. The lysis was apparent when detached cells were spun down, followed by replacement of the supernatant by fresh buffer, as this would result in an almost complete loss of β-galactosidase activity. This loss of activity indicates that the majority of the enzyme had been released from the cells. Such cell lysis also affects the quantification of the optical density. Indeed, when performing the experiments as described originally, even minute variations in sample handling, specifically during the vortexing and washing of the samples, led to highly variable results.

Instead, when exploiting the more robust normalization to protein content, experiments generated reproducible results. However, consistent with the results that we obtained from the single cell experiments, we found no difference in the expression of the reporter (p = 0. 14) between planktonic and sessile cells (Figure 1e). Thus, also with the original experimental approach, slightly adapted to increase reproducibility, we were unable to observe any surface-specific response of the CpxAR system.

## Discussion

Using both single-cell and population-level assays, we investigated surface sensing in *E. coli* via the CpxAR and RcsCDB systems. While we could confirm the strong induction of the Rcs system upon surface attachment, we could not identify activation of the Cpx system. The single-cell approach, involving microfluidics and fluorescence microscopy, showed a constant expression of the reporter gene following the switch from liquid culture to surface-attached growth. To exclude experimental difference as cause for this conflicting finding with an earlier report, we repeated the earlier presented population-level assay, where we also did not find any evidence for activation of the system.

One explanation for the disagreement between our results and the generally accepted view of CpxAR as a surface sensing system, could be that the original measurements on the population level had been confounded by technical factors, such as the cell lysis that we experienced upon detachment of cells from the beads. Such lysis is problematic both in terms of measurement of β-galactosidase activity and cell density quantification, which is necessary to normalize the measured activity. Also, later experiments by Shimizu *et al.* (25) may have suffered from the same technical issues, as their normalization to colony forming units is also expected to be highly sensitive to cell lysis. The previous finding that deletion of the *cpxR*, *cpxA* and *nlpE* genes abolished the response of the Cpx system to surface attachment (14), might instead be explained by the greatly reduced attachment of these mutants to hydrophobic surfaces. If the cells are weakly attached, as is the case in the mentioned deletion mutants (14), the lysis problems occurring at the detachment step may be alleviated, thereby removing the confounding effect.

An alternative explanation for the different observations could be that one of the many other, non-surface related factors induced the Cpx system in the previous studies. One possibility would be that the results had been affected by the presence of copper, which has a very strong effect on the Cpx system, even at low concentrations (Supplementary figure S1b). In fact, Cpx induction by copper strongly resembles the dynamics found in the original surface sensing experiments (14). Interestingly, the synthesis of the hydrophobic coating material, dimethyldichlorosilane, requires large quantities of copper (36) and possibly trace amounts might have been present in the experiments by Otto and Silhavy.

The original study, in which CpxAR had been established as a surface sensing system, is frequently cited to link the Cpx system and NlpE to adhesion and initiation of biofilm formation (e.g. (37–41, 33)). As shown in this work, the connection between the Cpx system and surface sensing needs to be reconsidered, to avoid incorrect interpretations of experimental findings.

## Materials and methods

### Bacterial strains and growth conditions

The MG1655 strain carrying transcriptional fluorescent reporters for the *yebE* gene (pPyebE-*gfp*) and *rcsA* gene (pPrcsA-*gfp*) were obtained from the *E. coli* promoter collection (42). The TR235 strain (MC4100 λRS88[*cpxR-lacZ*], (20)), which has a transcriptional reporter for the *cpxR* gene, was kindly provided by T.J. Silhavy. For microscopy experiments, the TR235 strain was transformed with the pPyebE-*gfp* plasmid. The MG1655∆*rcsB* strain was constructed by P1 phage transduction from the corresponding deletion strain in the Keio collection (43). After removal of the kanamycin resistance gene, the Δ*rcsB* strain was transformed with the pPrcsA-*gfp* plasmid.

Bacteria were grown at 37°C in an orbital shaker (300 rpm), in either M9 minimal medium supplemented with 0.4% glucose or LB medium. The medium was supplemented with 25 μg/ml kanamycin for the plasmid-carrying strains. Preparation of spent medium was done by spinning down bacterial cultures at 1000 g at 4°C and subsequent filtering of the supernatant through a 0.22 μm pore-size bottle-top filter made of PES. Spent medium was always taken from cultures at the same OD_600_ as the culture used for the experiment.

### Copper induction of Cpx and Rcs reporters

*E. coli* MG1655 with reporters for the Cpx (pPyebE-*gfp*) and Rcs (pPrcsA-*gfp*) systems were grown to mid exponential phase (OD 0.5 – 0.6) in M9 glucose medium without copper. The cultures were diluted 125 – 150-fold in fresh M9 glucose medium with or without 7 μM CuCl_2_, obtaining the same cell counts for all cultures, and measured at regular intervals by flow cytometry (BD Accuri C6 flow cytometer, BD Biosciences; medium flow rate, FSC-H-threshold 8000, SSC-H threshold 500). The fluorescence intensities in the GFP channel (FL-1) were normalized to the size of each cell, measured as the width. Each data point is the median of at least 36,000 cells.

### Preparation of hydrophobic surfaces

Cover glasses (Menzel-Gläser #1.5) were cleaned by a procedure adapted from (44): cover glasses were sonicated alternatingly in absolute ethanol and 2% Hellmanex III in ultrapure water; twice in each solvent, 30 minutes each, after which residual water was removed from the container by 10-minute sonication in acetone, followed by rinsing of the container with acetone twice. To apply the hydrophobic coating, the cover glasses were then incubated for 10 minutes with a 10% v/v solution of dimethyldichlorosilane in hexane, followed by extensive rinsing with absolute ethanol, in which the cover glasses were kept for no more than two weeks. The water contact angle (>85°) stayed constant during the two weeks, indicating the stability of the coating. The silanization of 0.5 mm diameter glass beads (Sigma-Aldrich G8772) was carried out in the same way, except for skipping the sonication steps, as the beads had been acid-washed by the manufacturer.

### Microfluidics

The microfluidic setup shown in Figure 1a was used. All components were prewarmed to 37°C. A 24 × 24 mm cover glass, either untreated (i.e. only rinsed with ethanol and ultrapure water) or hydrophobic (see above) as described in the main text, with a thin piece of PDMS around the edges to prevent leakage was placed in a custom-made metal holder. In the center of the cover glass, 5 μl of bacterial culture was pipetted and immediately covered by a 1.5 mm thick 10% polyacrylamide gel pad. This pad had been extensively washed after preparation and incubated for at least one hour in spent medium. The microfluidic setup was completed by a piece of PDMS containing a 2 × 10 mm channel that was placed on top of the pad. Using a plexiglass frame and bolts, the setup was tightened to the metal holder. Tubings (Cole-Parmer Microbore PTFE Tubing, EW-06417-11) were connected to the channel and spent medium was perfused at a flow rate of 24 μl/min throughout the experiment, for which a Harvard Apparatus syringe pump 11 Elite (#70-4505) was employed. Both the specimen and microscope were temperature-controlled to 37°C (Life Imaging Services, The Cube and The Box).

### Fluorescence microscopy

For image acquisition, a Nikon Eclipse Ti-E inverted microscope was used, with Nikon CFI Plan Apo Lambda DM 100X Oil objective, CoolLED pE-2 or Lumencor Aura illumination system (470 or 485 nm LED, respectively, for excitation of GFP) and Andor iXon 897 EM-CCD camera. The following filters were employed: excitation filter bandpass 470/40 nm, dichroic mirror 495 nm and emission filter 525/50 nm (AHF Analysentechnik F46-470). Focus was maintained by Nikon’s PFS3 system. Acquisition was started within 10 min (generally ~7 min) after the bacteria had been introduced in the microfluidic system and every 10 min phase contrast images and GFP signal were acquired at multiple positions. The microscope was controlled by NIS Elements v4.51 software.

### Image analysis

Image segmentation was semi-automated and handled by the ImageJ (45) plugin MicrobeJ (46), or in-house software, followed by manual inspection and correction. The detected cells were further analyzed in Matlab (R2014a, MathWorks Inc.), where the identified ROIs were applied to background-corrected GFP images, obtained by subtracting the signal intensity of images without any bacteria, and application of a correction for uneven illumination as quantified from the background.

### Population-level assay

The assay to study CpxR activity on the population level in bead-attached cells was carried out essentially as described (14). To plastic tubes containing 3 g of freshly prepared hydrophobic beads (prepared as described above in ‘Preparation of hydrophobic surfaces’), 1 ml of OD_600_ 2.0 culture of TR235 in LB was added and incubated at 37°C. After 1 h, unattached cells were aspirated using a syringe with needle and discarded. Attached cells were then detached by addition of 1 ml Z-buffer containing 0.04% SDS, vortexing for 30 s and aspirating with a syringe with needle. Cells were lysed by addition of three drops of chloroform and vortexing for 15 s. As planktonic controls, bacteria incubated in tubes without beads were used. These control cells were spun down, resuspended in Z-buffer with 0.04% SDS and three drops of chloroform, and vortexed for 15 s. From all samples, 50 μl was set aside for determination of total protein content, and the remainder was used for the β-galactosidase assay.

### β-galactosidase assay

The assay was carried out essentially as originally described by Miller (47). All samples were incubated at 28°C and the reaction was started by addition of 200 μl 4 mg/ml ONPG (Sigma-Aldrich #N1127). Reactions that had turned yellow upon visual inspection were stopped by mixing with 500 μl 1 M Na_2_CO_3_. The samples were spun down and the absorption at 420 nm of the supernatant was measured. The β-galactosidase activity was calculated as (1000 • A_420_) / (TP • t), where ‘TP’ is the total protein concentration in relative units and ‘t’ is the duration of the reaction in minutes.

### Determination of total protein content

The protein content in the β-galactosidase samples was determined from the band intensities on a silver-stained 10% SDS-PAGE gel. The gel was stained according to the procedure provided with the kit (Pierce Silver Stain Kit, #24612). The stained gel was imaged with a Fujifilm LAS-3000. In ImageJ (45), the background signal was determined and the intensities of the bands in each gel lane were integrated. After background correction, the total intensity was normalized to a control sample, which was included on multiple gels.

## Acknowledgements

We would like to thank Thomas J. Silhavy for kindly providing the TR235 strain. This work was financed by the Netherlands Organisation for Scientific Research (NWO) through a VIDI grant to MH (project number 864.11.001).

---

**Supplementary figure 1:**
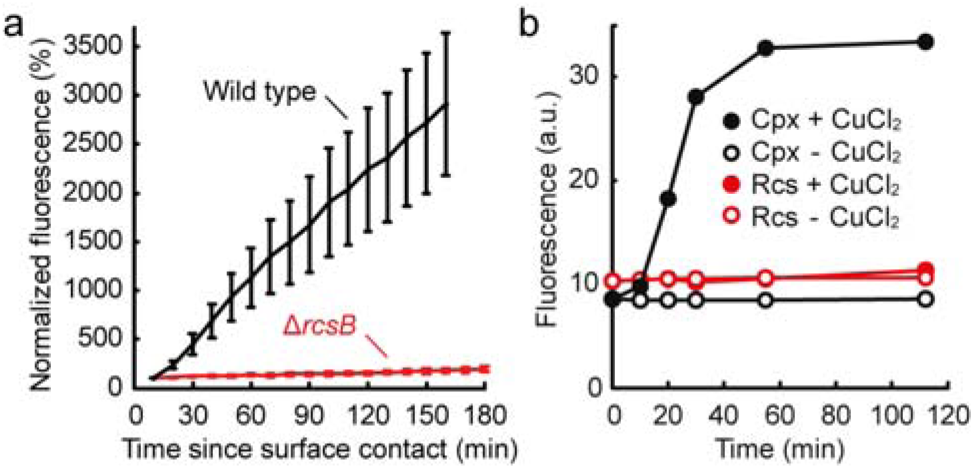
(a) Comparison of GFP expression from the Rcs-regulated *rcsA* promoter in wild type (black; n = 46; 2 independent experiments) and Δ*rcsB* cells (red; n = 40; 2 independent experiments) on untreated cover glasses, in the microfluidic device with flow of spent M9 glucose medium. The fluorescence intensity of each cell at the first time point is set to 100%. Error bars show 95% confidence intervals. (b) Effect of copper chloride on the reporters for the Cpx (pPyebE-*gfp*) and Rcs (pPrcsA-*gfp*) systems. Exponential phase M9 glucose cultures were diluted in fresh M9 glucose medium with or without 7 μM CuCl_2_ and measured at regular intervals by flow cytometry. The fluorescence intensities were normalized to the size of each cell and shown here as the median of at least 36,000 cells.

